## Supplementary figures for "Mother-Infant Gut Viruses and their Bacterial Hosts: Transmission Patterns and Dynamics during Pregnancy and Early Life"

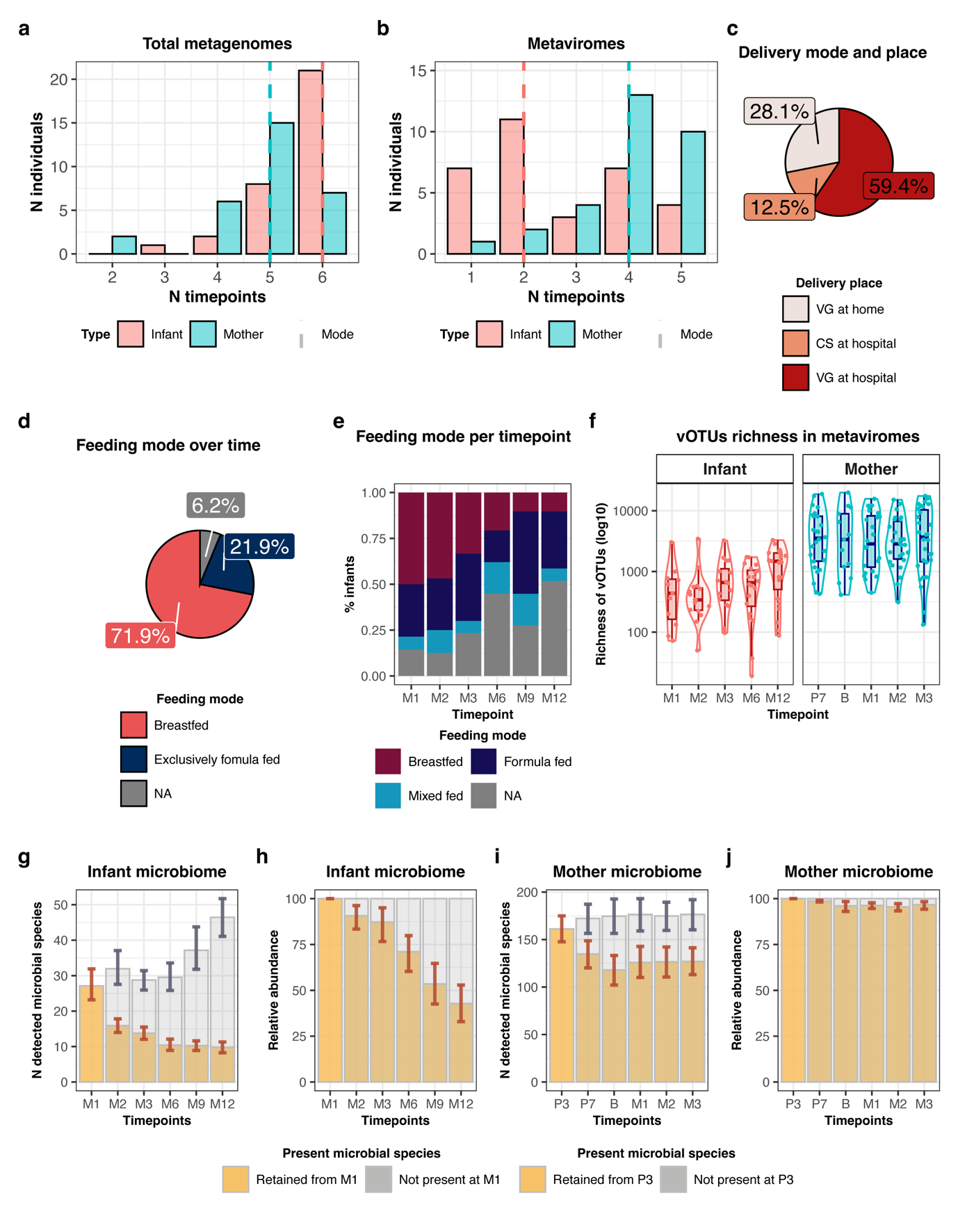
 **Supplementary Figure 1: Study population, vOTU richness in metaviromes, and microbiome stability.**

**a**, **b,** Number of timepoints in **a**, total metagenomes and **b**, metaviromes for infants (pink) and mothers (cyan). Dashed lines of respective colors depict the mode for infants and mothers separately. **c**, Pie chart showing the distribution of infants by delivery mode and place of delivery (home vs hospital). CS: Cesarean section, VG: vaginal delivery. **d**, Pie chart showing the distribution of the infant feeding mode across all timepoints. **e,** Stacked bar plot showing the infant feeding mode per timepoint. Y-axis depicts the percentage of infants within each feeding mode category at the given timepoint. **f,** vOTUs richness in infant and maternal metaviromes at different timepoints. **g,** Number and **h,** relative abundance of bacterial species retained from month 1 in infants at months 2, 3, 6, 9 and 12 after birth. **I,** Number and **j,** relative abundance of bacterial species retained from the 3rd month of pregnancy in mothers to the 7th month of pregnancy, delivery, and months 1, 2, 3 after delivery.

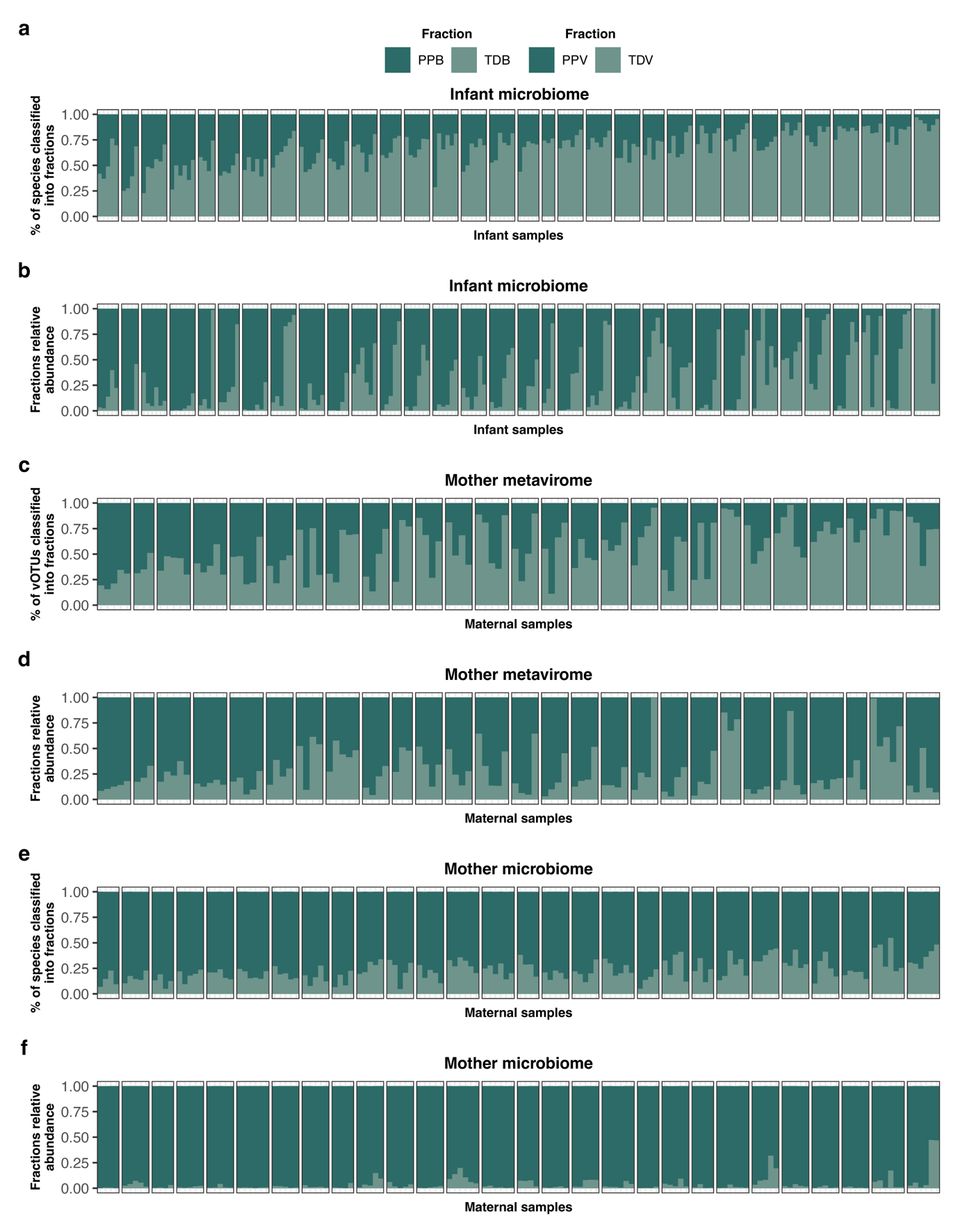

**Supplementary Figure 2: Size and abundance of personal biome fractions.**

**a,** Percentage of bacterial species classified into biome fractions defined based on the species prevalence in the infant microbiome. Personal persistent bacteriomes (PPBs) are those present in >= 75% of individual samples and shown in dark green. Transiently detected bacteriomes (TDBs) are those present in less than 75% of the samples of an individual and are shown in light green. Each facet depicts one timepoint of an infant. **b,** The relative abundance of biome fractions in the infant microbiome (PPB and TDB). **c,** Percentage of vOTUs classified into biome fractions defined based on the vOTUs prevalence in the maternal virome. Personal persistent viromes (PPVs) are those present in >= 75% of individual samples and shown in dark green. Transiently detected viruses (TDVs) are those present in less than 75% of the samples of an individual and shown in light green. **d,** The relative abundance of biome fractions in maternal viromes (PPV, TDV). **e,** Percentage of bacterial species classified into biome fractions is defined based on the prevalence of bacterial species in the maternal microbiome (PPB, TDB). **f,** The relative abundance of biome fractions in maternal microbiomes (PPB, TDB).

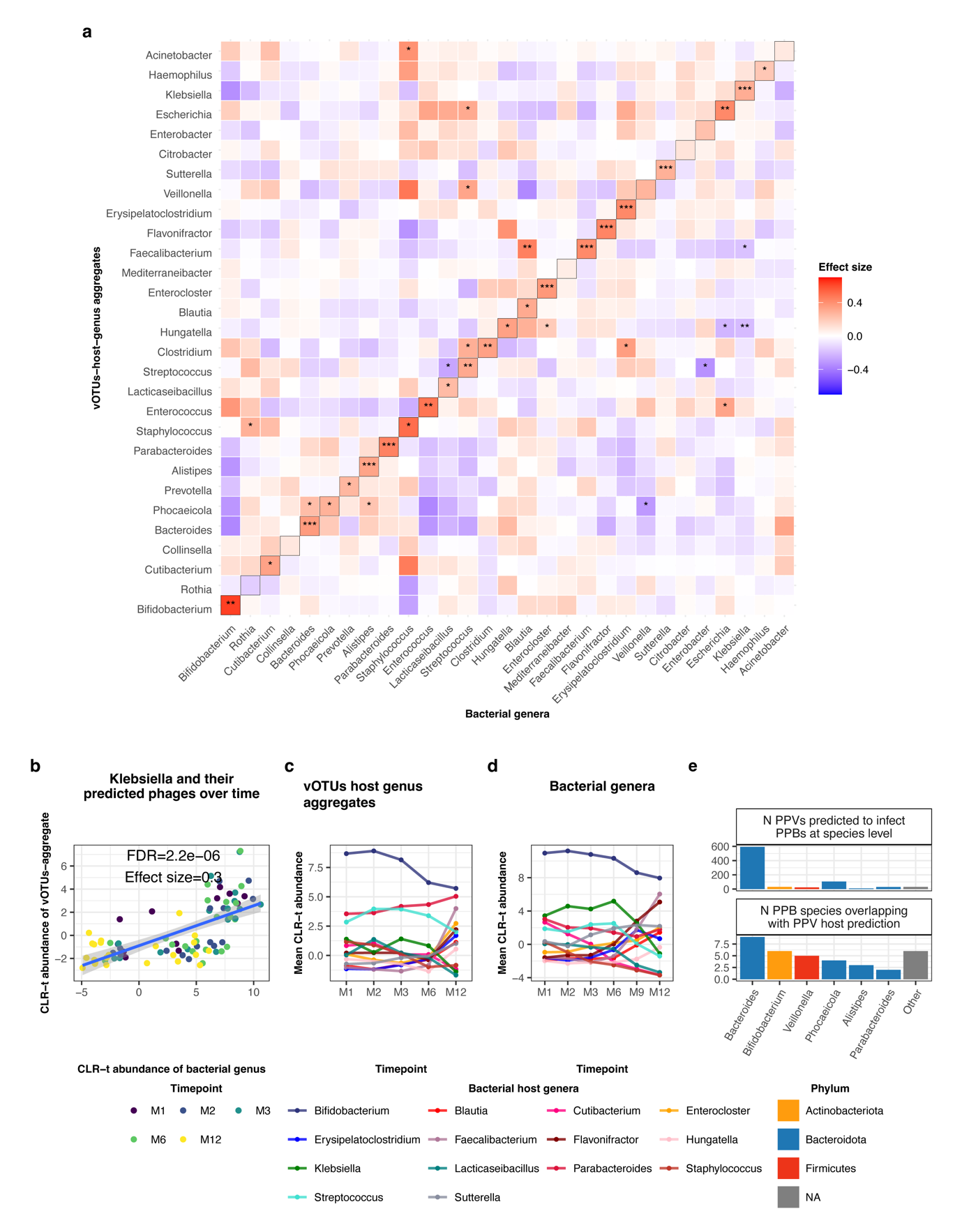

**Supplementary Figure 3: Correlation between infant vOTU-host-genus aggregates and bacterial genera and their dynamics in infants.**

**a,** Heatmap showing the associations between vOTUs, aggregated based on their hosts at the genus level and their predicted bacterial genera. The color represents the effect size, and the stars indicate FDR significance. *** indicates FDR-values < 0.001, ** indicates FDR-values <0.01, and * indicates FDR-values <0.05. **b,** A scatterplot showing the association of the centered log ratio (CLR) transformed vOTU abundances, aggregated based on their predicted host *Klebsiella*, on the Y-axis versus the CLR-transformed abundance of the bacterium *Klebsiell*a on the X-axis. **c,** Dynamics of vOTU host aggregates over time in infants (only vOTU aggregates prevalent in >20% of infant metaviromes and significantly associated with their predicted host at the genus level and time are shown). **d,** Dynamics of the bacterial genera predicted as hosts for vOTU aggregates over time in infants. **e,** Bar plot showing the number of PPVs predicted to infect PPBs at the species level (top) and the number of overlapping PPVs and PPBs by the PPVs’ host prediction (bottom) within the most prevalent genera.

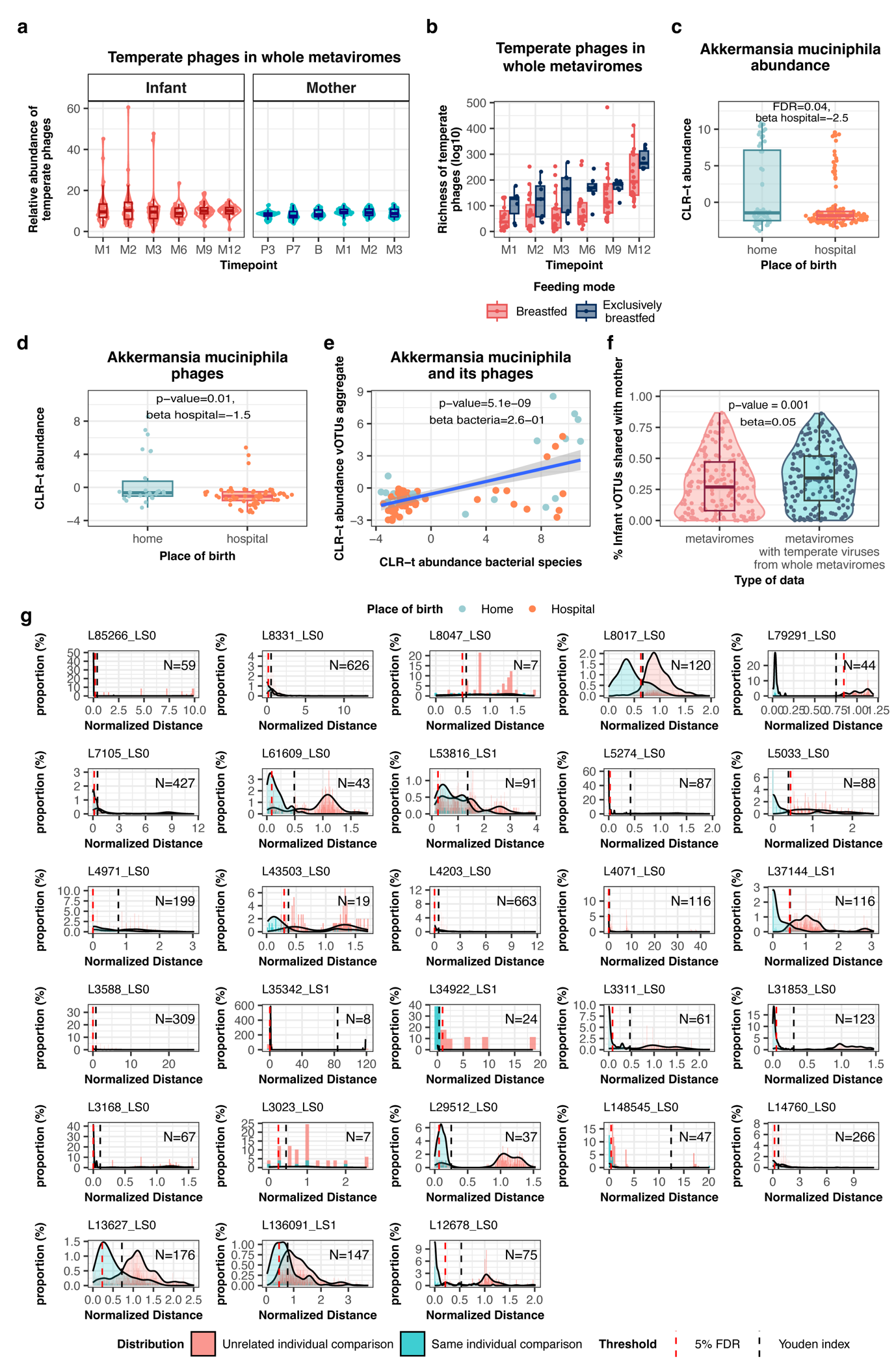

**Supplementary Figure 4: Temperate phages in whole metaviromes, gut virome phenotype association and within-individual virus strain variation.**

**a,** Relative abundance of temperate bacteriophages in infants and mothers in whole metaviromes over time. **b,** Boxplots showing the richness of temperate bacteriophages in whole metaviromes in infants, with colors representing the feeding mode. **c,** Boxplots showing the difference in CLR-transformed abundance of *Akkermansia muciniphila* between infants born at home versus those born in a hospital (all timepoints are pooled for the figure). Colors represent different places of birth. **d,** Boxplots showing the difference in CLR-transformed aggregated relative abundance of A. muciniphila phages between infants born at home versus those born in a hospital. **e,** Correlation between CLR-transformed abundance of *A. muciniphila* and CLR-transformed aggregated abundance of its phages. **f,** Violin plots and boxplots showing the increase in sharedness of infant vOTUs to maternal vOTUs detected in metaviromes upon the inclusion of prophages detected in whole metaviromes. **g,** Distribution curves of virus genetic distances between strains detected in longitudinal samples of the same individual (blue) and between strains detected in samples of unrelated individuals (red) for 28 viruses that showed lower genetic distance between strains detected in related mother-infant pairs compared to unrelated mother-infant pairs. X-axis depicts the normalized by median genetic distance attributed to each virus. Strain identity thresholds were set as the Youden’s index (black dashed line) or as an empirical FDR (the 5th percentile) of the unrelated individual comparisons (red dashed line) when the first was above 5%. The N in each histogram corresponds to the number of same-individual comparisons.

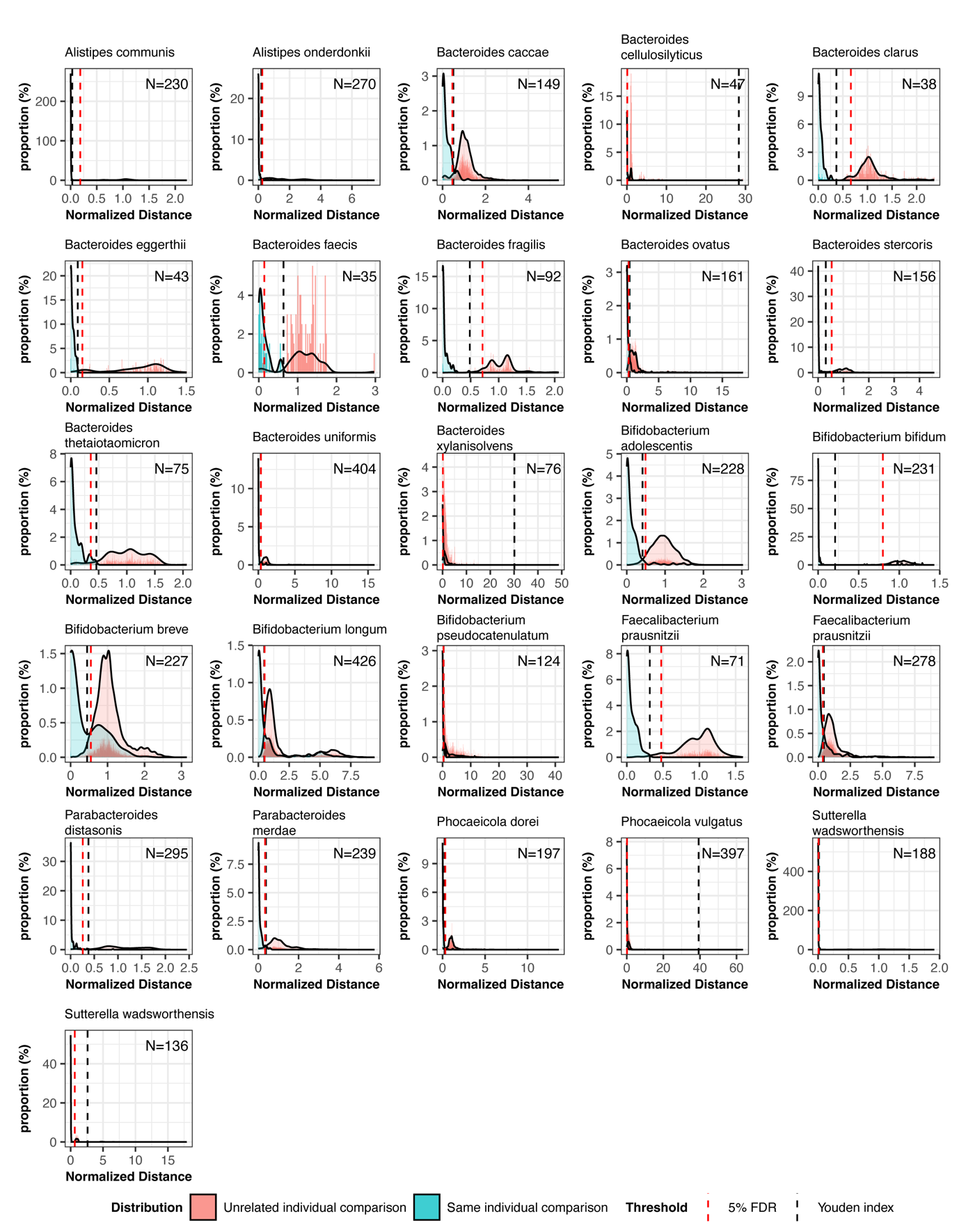

**Supplementary Figure 5:** **Within-individual bacterial strain variation**

Distribution curves of bacterial genetic distances between strains detected in longitudinal samples of the same individual (blue) and between strains detected in samples of unrelated individuals (red) for 26 bacterial strains that were predicted to be hosts for transmitted viruses and showed lower genetic distance between strains detected in related mother-infant pairs compared to unrelated mother-infant pairs. X-axis depicts the normalized by median genetic distance attributed to each bacterial strain. Strain identity thresholds were set as the Youden’s index (black dashed line) or as an empirical FDR (the 5th percentile) of the unrelated individual comparisons (red dashed line) when the first was above 5%. The N in each histogram corresponds to the number of same-individual comparisons.

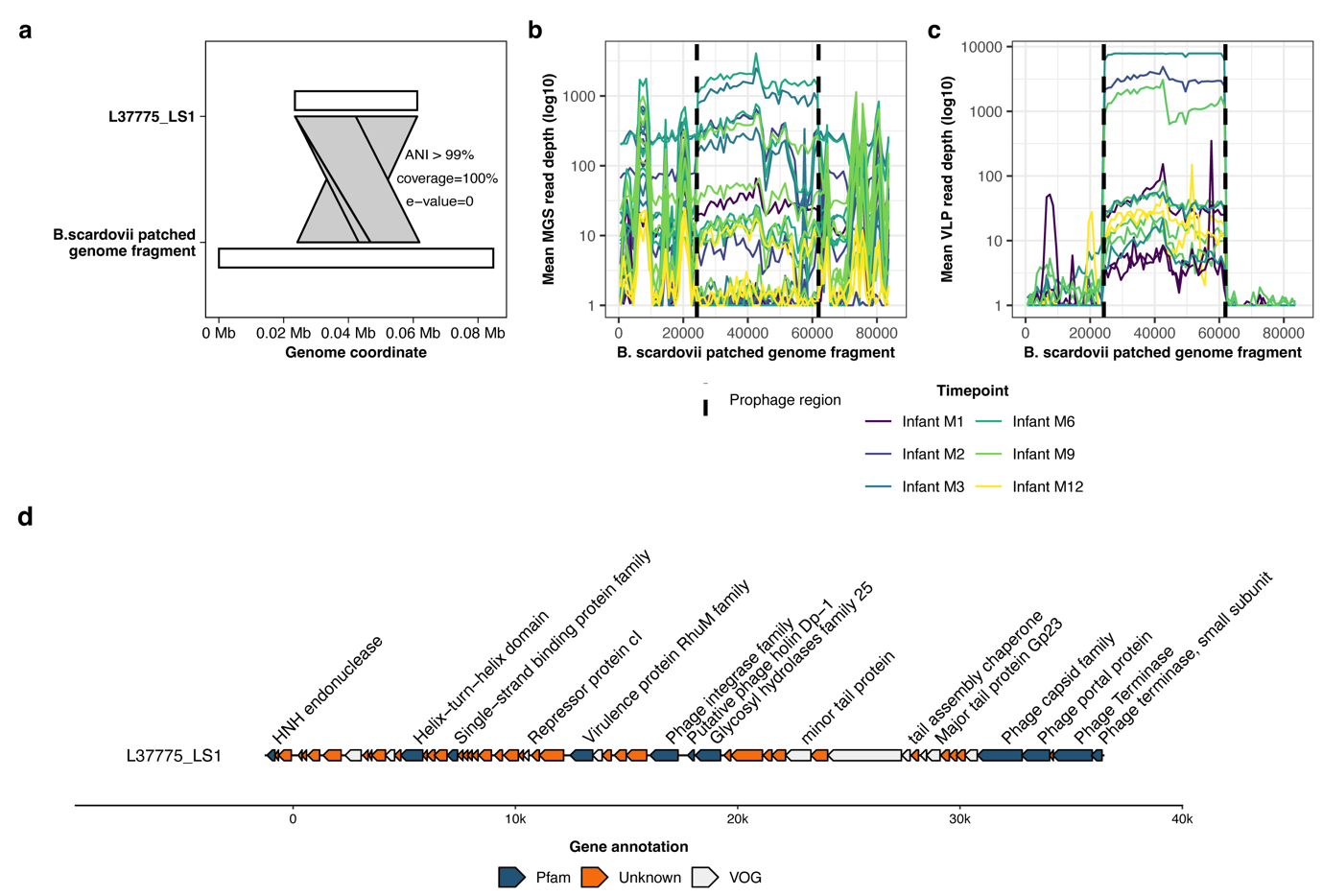

**Supplementary Figure 6: Example of virus-host cotransmission not rooted in the maternal gut: a temperate phage of *Bifidobacterium scardovii***

**a,** Synteny plot of L37775_LS1 genome sequence mapping to the *Bifidobacterium scardovii* patched genome fragment. X-axis indicates the genome coordinates (in megabases). Lines connecting the L37775_LS1 and *B. scardovii* indicate the prophage insertion region in the bacterium genome fragment. **b,** Depth of the *B. scardovii* patched genome fragment coverage by sequencing reads from total metagenomes that were positive for *B. scardovii,* and **c**, metaviromes that were positive for L37775_LS1. Each color line corresponds to an infant sample and represents mean depth in a 1,001-nt sliding window. Line colors represent samples from different infant timepoints. X-axis depicts the genome coordinate in base pairs. Dashed lines indicate the prophage insertion region. **d,** Genome organization of L37775_LS1. X-axis depicts the genome coordinates in base pairs. Every predicted protein is represented by a polygon, and its orientation indicates the location of the predicted protein at the positive (right-orientation) or negative (left-orientation) strands. Colors indicate the source of protein annotation: Pfam (dark green), VOG (white), and proteins with no functional annotations are shown in orange. Hypothetical and annotated proteins of unknown function are shown in respective colors without text annotation.
